# Automated phenotyping of mosquito larvae enables high-throughput screening for novel larvicides and offers potential for smartphone-based detection of larval insecticide resistance

**DOI:** 10.1101/2020.07.20.211946

**Authors:** Steven D. Buckingham, Frederick A. Partridge, Beth C. Poulton, Ben Miller, Rachel A. McKendry, Gareth J Lycett, David B. Sattelle

## Abstract

Pyrethroid-impregnated nets have contributed significantly to halving the burden of malaria but resistance threatens their future efficacy and the pipeline of new insecticides is short. Here we report that an invertebrate automated phenotyping platform (INVAPP), combined with the algorithm Paragon, provides a robust system for measuring larval motility in *Anopheles gambiae (and An. coluzzi)* as well as *Aedes aegypti* with the capacity for high-throughput screening for new larvicides. By this means, we reliably quantified both time- and concentration-dependent actions of chemical insecticides faster than using the WHO standard larval assay. We illustrate the effectiveness of the system using an established larvicide (temephos) and demonstrate its capacity for library-scale chemical screening using the Medicines for Malaria Venture (MMV) Pathogen Box library. As a proof-of-principle, this library screen identified a compound, subsequently confirmed to be tolfenpyrad, as an effective larvicide. We have also used the INVAPP / Paragon system to compare responses in larvae derived from WHO classified deltamethrin resistant and sensitive mosquitoes. We show how this approach to monitoring larval response to insecticides can be adapted for use with a smartphone camera application and therefore has potential for further development as a simple portable field-assay with associated real-time, geo-located information to identify hotspots.

**Author summary:** We have developed an automated platform for recording the motility of mosquito larvae and applied it to larvae of a mosquito vector of malaria and a mosquito vector of dengue, Zika, yellow fever and other human diseases. The platform facilitates high-throughput, chemical screening for new compounds to control mosquito larvae and also detects differences in response in larval progeny from pryethroid resistant adult mosquitoes. Pyrethroid-impregnated bednets have helped to halve the deaths from malaria in recent years but pyrethroid resistance is an important threat to this progress. Our approach assays insecticide actions faster than the current WHO standard test and we show that it can be adapted for use with a smartphone, which offers the prospect of a future field assay to monitor developing resistance associated with behavioural response with the added benefit of precise satellite-based location.

## Introduction

Mosquito-borne pathogens kill more than one million people every year, while severely debilitating diseases including malaria, dengue, Zika, West Nile virus, chikungunya, yellow fever and Japanese encephalitis, transmitted by these vectors cause suffering for hundreds of millions. Successful attempts to control the incidence of such diseases have mainly targeted mosquito population size and hence infective biting frequency [1]. As a result, the burden of malaria has been halved in the period 2000-2015, mostly due to the use of insecticide-treated nets (ITNs) containing pyrethroids, which target the adult disease-transmitting Anopheles mosquitoes that tend to feed nocturnally in homes [2]. However, the future efficacy of ITNs to control malaria is threatened by growing mosquito resistance to the available pool of insecticides [3] and by an increase in the prevalence of Anopheles mosquitoes that blood-feed outdoors [4].

ITNs are of limited use against viral pathogens transmitted by *Aedes aegypti* mosquitoes, such as dengue, Zika, yellow fever and chikungunya, since the adult vectors favour feeding at dawn and dusk. The ineffectiveness of ITNs against Aedes and the absence of drugs or vaccines for most vector-borne viruses means that a mainstay of viral disease control has been larval source management (LSM) [5]. This integrated approach targets aquatic egg-laying sites with the aim of reducing the number of immature forms, leading in turn to fewer biting adults. LSM can incorporate community environmental measures to limit temporary aquatic breeding sites in and around homes, as well as the application of larvicidal agents, such as temephos, to important, permanent water sources. These traditional LSM methods are effective where the breeding sites that produce the majority of adults can be readily identified and treated accordingly [6]. Alternative methods, attempting to overcome this limitation, involve the dusting of resting female mosquitoes with highly potent larval growth inhibitors. As a result, growth inhibitors are delivered to oviposition sites at doses proportional to the frequency of visits. By this means, the most important breeding sites are targeted [7].

For malaria control, there is growing awareness that LSM may have a more prominent role to play under certain situations, complementing adult-targeted insecticide interventions. Examples include: targeting zones where the prominent vectors rest and feed outdoors; tackling transmission foci, where saturating ITN usage is having limited effect on disease transmission; deployment in areas where adults are resistant to all known insecticides; treatment of urban areas in which water sources that suit Anopheline development may be more readily identified [8]. Technologies, such as drones and mapping have been combined to improve identification and prioritisation of habitats in rural areas [9,10]. There may be additional opportunities to control multiple larval mosquito species simultaneously that will affect both malaria and viral disease transmission [6].

Larvicides in current use include surface oils and films that suffocate immature forms, bacteria (including *Bacillus thuringiensis* that produces toxic protein crystals that target the larval midgut), insect growth regulators (including pyriproxyfen that inhibits adult emergence) and synthetic chemicals (such as chlorpyrifos, pirimiphos-methyl and temephos) that target the insect nervous system [6].

Standardized methods to examine relative larval susceptibility to compounds are necessary to maintain the effective deployment of current larvicides by identifying the emergence of resistance in mosquito populations [11]. Moreover, a high-throughput method to screen larvicides would offer the possibility to examine large chemical libraries for new classes of compounds. Currently the World Health Organisation issues guidelines for larvicidal toxicity assays [12], which consist of exposing larvae in small cups to increasing doses of chemical and manual observation of mosquito killing. In such assays, larval death is defined by the moribund appearance of larvae and their failure to respond to tapping of the cup. This is labour-intensive and the end-point can be difficult to assign unambiguously.

We have developed an automated phenotyping assay based on larval motility, which is simple to deploy, provides a fast readout and, being multi-well plate based, offers much needed high-throughput screening capabilities. Our system uses an <u>INV</u>ertebrate <u>A</u>utomated <u>P</u>henotyping <u>P</u>latform (INVAPP) previously described for monitoring nematode motility in conjunction with the Paragon algorithm to estimate motility from moving images of larvae [13]. We illustrate the capability of our system by (a) quantifying the actions of the larvicide temephos on larval motility, (b) screening a chemical library, which showed larvicidal actions of tolfenpyrad and (c) detecting differential responses in larval progeny from deltamethrin resistant and susceptible *Anopheles coluzzii* and *Aedes aegypti* mosquitos. We also demonstrate the potential for its further development as a smartphone-based field assay, for rapid detection of behavioural change related to pyrethroid resistance, using the inbuilt camera to track larval motility.

## Methods

### Mosquito rearing and strains

The mosquito strains used are established colonies which have been characterised by WHO protocols as either deltamethrin sensitive or resistant in female adults. They have been in lab culture for 2 - 20 years and were not infected with any known pathogen. Resistant (Cayman) and sensitive (New Orleans) *Ae. aegypti* strains and resistant *An. coluzzii* (Tiassale) and sensitive *An. gambiae* (G3) strains were studied. The resistant lines were obtained from LITE (https://lite.lstmed.ac.uk/lite-facilities/lite-insectaries), which regularly (approx. every 3 months) select adults with permethrin/deltamethrin. Cayman show between 20-50 % mortality and Tiassale 10-20% moratality to deltamethrin in standard WHO bioassays. These strains are not characterised routinely for larval resistance to temephos and deltamethrin. Previous assays on the Cayman larvae indicated a 1.64 resistance ratio compared to the New Orleans strain (Harris 2010). The mosquito batches maintained at LSTM and provided to UCL were not further selected prior to despatch. It should be noted that there exists a considerable degree of genetic variation in Anopheline mosquitoes especially in the species complex *Anopheles gambiae sensu lato*, which is widespread in sub-Saharan Africa. *Anopheles gambiae* and *Anopheles coluzzii* are morphologically indistinguishable species and were for many years considered as distinct molecular forms (S and M respectively) of the same species. The G3 strain we here deploy has always been called *Anopheles gambiae* but is in fact a hybrid of molecular forms. Pure *An. coluzzii* strains can be found in Tiassale, Cote D’Ivore, but the lab strain Tiasalle 13 deployed here is also a hybrid.

### Larval hatching and maintenance

#### Anopheles

Anopheles eggs laid day 1 evening or day 2 morning were shipped securely from Liverpool School of Tropical Medicine (LSTM) to University College London (UCL) on filter paper ensuring that they remain moist throughout the journey. At UCL on day 3, eggs were washed into a shallow dish containing deoxygenated (pre-boiled and cooled) water containing 0.001% pond guardian tonic salt (Blagdon) to a depth of about 2.5 cm. The eggs and newly-hatched larvae were maintained at 25°C. Larvae were fed 1/3 pellet of cat food per dish and tested on days 5-6.

#### Aedes aegypti

*Ae. aegypti* egg papers were shipped securely to UCL from LSTM and stored at room temperature for up to 1 month. They were hatched over 1-2 days by placing the egg papers in a tray of deoxygenated (pre-boiled and cooled) tap water to which a crushed yeast tablet (Holland and Barrett) had been added. Larvae were tested on days 3-4.

#### Harvesting larvae and transfer to 96-well plates

Experiments were performed on larvae 1-2 days post hatching and thus a mix of first and early 2^nd^ instar larvae. The water in which larvae were swimming was passed through a 100 μm Nylon mesh cell-strainer (Fisher Scientific) to concentrate the larvae. This concentrated suspension of larvae was diluted until 100 μL of suspension contained 5-10 larvae. A 100 μL aliquot of this suspension was added using a standard Gilson pipette to each well of a 96-well plate, with the tip of the pipette being cut back to reduce damage to the larvae. Subsequently, 100 μL of the compound under test was added (dissolved in water from a 10^−2^; M DMSO stock to yield the required final concentration of 10^−4^M). Wells containing DMSO alone diluted to appropriate concentrations served as controls.

#### Filming larvae using INVAPP and analysis of motility using Paragon

Larvae were filmed using INVAPP, a device for monitoring invertebrate motility that is well suited to high-throughput chemical screening and has been deployed for the study of nematode motility (11). INVAPP consists of an Andor Neo “sCMOS” camera (2560 x 2160 resolution, maximum 100 frames per second frame rate) fitted with a Pentax YF3528 line-scan lens. The camera is mounted beneath a microplate supporting platform illuminated from above by an LED array panel. An acrylic diffuser ensures that the field of view of the camera is evenly lit (Figure 1).

**Figure 1.**
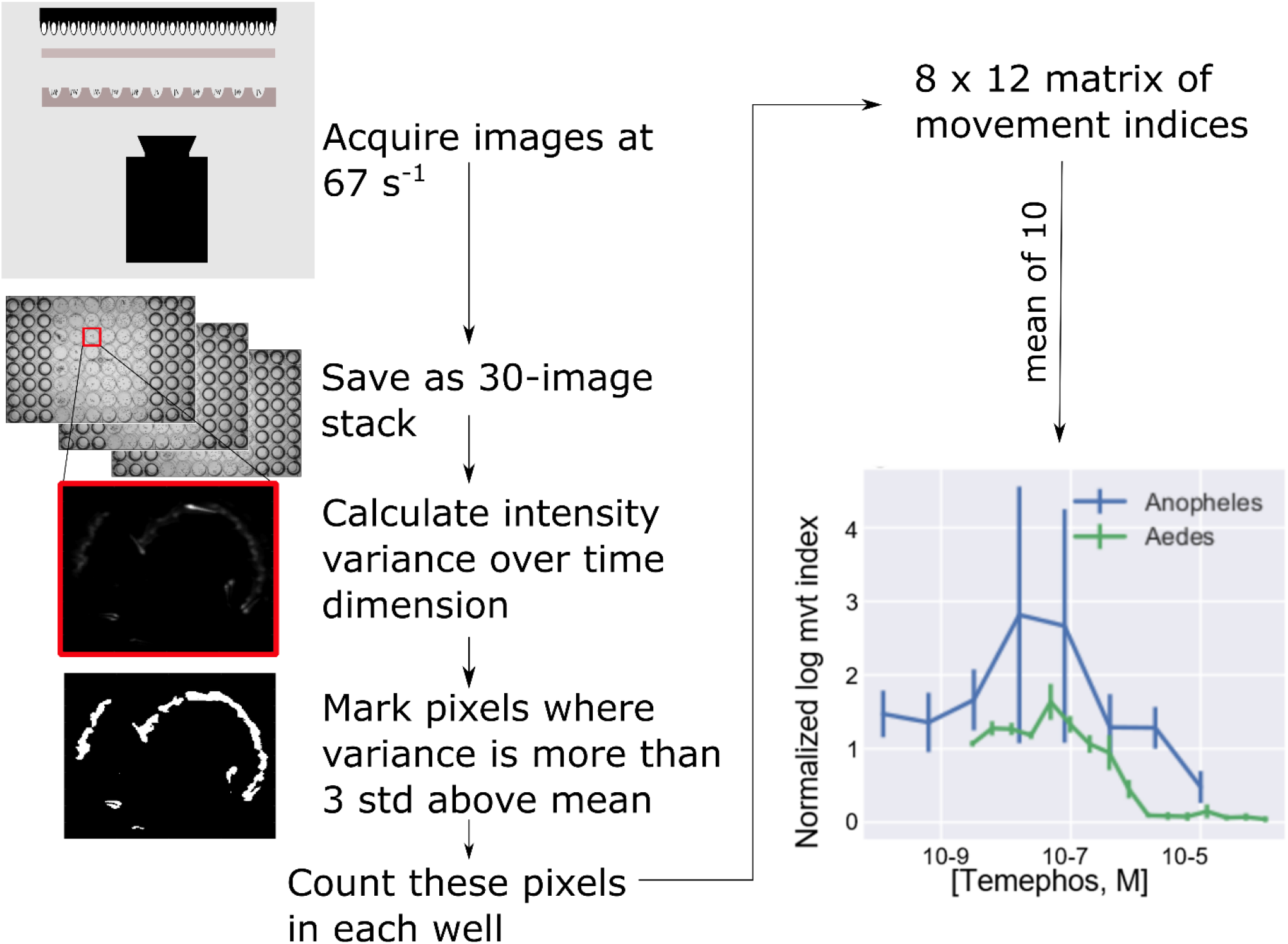
The procedure for automating the analysis of mosquito larval swimming. In each trial, 30 images are acquired at 10 ms intervals and stored for later offline analysis. An index of the amount of movement is obtained by measuring the variance for each pixel over time. Pixels for which the variance more than 3 standard deviations from the mean variance are scored as 1, the remainder as 0. The movement index for each well is taken as the sum of these scores for that well. The output of the algorithm for quantifying movement plotted against the concentration of temephos (bottom right), a larvicide commonly used in the control of mosquitoes, is shown at the end of the pipeline. Alongside this similar data for larvae of *An. gambiae* are shown. A concentration-dependent inhibition of movement is seen in studies on larvae of *An. gambiae* and *Ae. aegypti*.

Images (total = 30) were acquired every 10 ms. This was repeated at approximately 5 s intervals until 5 or 10 series of image sequences were obtained. Acquiring images at this rate detected the slow, drifting movements associated with filter feeding as well as rapid “jumping” that larvae undergo at sporadic intervals. For every plate, in addition to filming at selected time points, readings were collected immediately prior to addition of chemicals. Image sequences were stored offline for later analysis. This was performed at least three times in separate weeks with each of the three replicates consitituting one data point in the analysis (i.e. at least three biological replicates).

#### Paragon algorithm

The analysis uses an implementation of the Paragon algorithm [13, https://github.com/fpartridge/invapp-paragon] and is described in outline below:

1. Read frames 1, 11 and 21 of the image sequence into a *n×m×*3 array where *n, m* are the height (approximately 700 pixels) and width (approximately 950 pixels) of the image sequences.
2. C alculate the variance in the time dimension
3. Identify the pixels for which the variance exceeds the mean by 3 standard deviations:

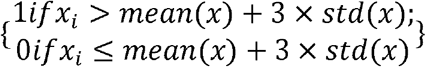
4. For each well, count the number of pixels of value 1 to produce a Movement Index
5. Normalize the movement index for all time points by dividing it by the value obtained for movies acquired immediately before adding compounds.

The procedure was performed three times and the median value is taken as the output of the screen. Image processing and inferencing was performed with Python scripts utilizing the following publicly available software libraries: Numpy, Pandas, Scikit-Image and Statsmodels. Matplotlib and Seaborn software libraries were employed for plotting.

Filming a plate requires 5 (in some cases 10) image sequences to be made each with 30 frames at 10 ms intervals, requiring a total of about 1 s for each image sequence. Analyzing one movie takes approximately 3 s on a desktop computer. The speed of recording and analysis permits library-scale screening of re-profiled and novel, candidate insecticide chemistry.

Estimation of pIC50

The pIC50 were estimated by fitting the concentration and the movement index to the expression:

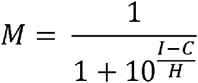

Where M is the predicted normalized movement index (the index measured after the application of compound divided by that measured before), I is the IC50, C is the concentration of the compound and H is the slope. Fitting was done by the least squares method using the Python “Scipy” package. The pIC50 is the −log10 of the IC50.

#### Chemicals

Temephos, an organophosphate anti-cholinesterase and established mosquito larvicide, was obtained from Sigma Aldrich (Item 31526-250MG). Deltamethrin (DM) is a pyrethroid insecticide used in bednets for which acts on insect sodium channels [14–16] and was obtained from Chem Service (Item N-11579). This particular pyrethroid was selected for study as the strains used had previously been characterised for deltamethrin resistance in the adult stage, and we were keen to examine if resistance could also be detected during larval stages using the INVAPP / Paragon system. Temephos and deltamethrin were prepared as stock solutions (1 x 10^−2^ M in DMSO) then diluted to the required final concentrations in water.

The Medicines for Malaria Venture Pathogen box library contains 400 drug-like molecules targeting a wide range of neglected tropical diseases. We used this to explore the potential of library-scale drug screening using INVAPP / Paragon.

#### Smartphone-based detection of motility

The smartphone application software was written using Android Studio with the OpenCV 3.3.0 library to implement a camera, handling the RAW pixel values in real time. The smartphone used was an LG G4 (16 MP, f/1.8, 28mm). After adding the larvae to wells of a 96 well plate and leaving to settle, recording is initiated with the phone camera pointing at the well plate. The application then automatically initialises two 1D arrays for the real-time mean and variances of each pixel. On each camera frame, the raw pixel values are extracted as a 1D byte array and converted to a float array. The mean and variance matrices are updated with the following formulae:

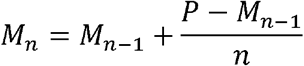

where, M is the mean matrix, P is the pixel value matrix and n is the number of frames.

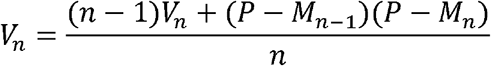

where V is the variance matrix.

After 60 frames, the mean and variance matrices are exported as a text file. This measurement is repeated every 10 s. The frame-rate, and therefore sensitivity, depends on the resolution of the image. Here, 320×240 images are used, giving around 30 frames per s, and therefore 2 s measurements every 10 s.

The variance matrix gives high values for areas of the frame where there is high motility (i.e in the wells with high larval motility), and low values (noise) between wells. This matrix can then be segmented into regions of interest (wells with larvae). This is a flexible approach, allowing measurement of a whole plate, or parts of a plate, depending on the framing.

At t=200 s, deltamethrin or control (water) was added to each well, and the variance tracked over time as described. After segmenting the regions of in interest, the variance at each time point for each well is calculated by taking the mean of the variance in the well. These are then plotted over time, showing the variance change with time. The resulting time-variance plots are fitted to exponential a decay function (Mathematica 11.3):

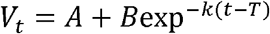

where *V* is the variance, *A* is the baseline variance, *B* is the maximum variance due to the larvae, *k* is the variance decay rate, and *T* is the time when the variance begins to decrease.

The fitted values for *k* are used to distinguish between susceptible and resistant larvae.

#### WHO Larval Assay

The WHO larval assay was published in 2005 with the aim of standardizing testing of compounds for their larvicidal actions on mosquitoes. It provides a detailed protocol for quantifying these toxic actions of insecticides on mosquito larvae in the field and in the laboratory and is available from https://apps.who.int/iris/bitstream/handle/10665/69101/WHO_CDS_WHOPES_GCDPP_2005.13.pdf. Briefly, a quarter yeast tablet (Holland and Barrett) was added to 200 ml of dH_2_O containing 0.001% pond salt (Blagdon), prior to addition of 25 third-instar *Ae. aegypti* (New Orleans or Cayman) larvae per cup. The appropriate volume of deltamethrin stock (dissolved in acetone) was then added to achieve the desired concentrations (5 x 10^−10^ – 1 x 10^−6^ M respectively). The assays were performed in triplicate and repeated three times on different larval batches. Mortality was assessed visually after 24 h continuous exposure.

## Results

### Using the INVAPP system to study the actions of the larvicide temephos on larvae of Aedes aegypti

Mosquito larvae swim [17,18], jump [19] and rest [18]. Filming at 1 frame every 10 miliseconds ensures that all types of larval movements are captured, but in accordance with the system’s goal of producing a rapid and simple index of larval activity these activities are not distinguished – the system simply “decides” how much area in the image has seen movement of any kind. Temephos, a widely used larvicide targeting *Ae. aegypti*, was tested on susceptible larvae of this species. A concentration-dependent reduction in motility was detected after 240 min exposure to this larvicide (Fig. 1, bottom right panel). Thus, the INVAPP / Paragon system enables robust, automated detection of larval motility and the severe motility impairment resulting from exposure to the larvicide temephos.

### Screening a chemical library for compounds with larvicidal activity

Given the system’s ability to provide a monitor of impaired motility in the presence of an established larvicide, we tested the system using a small chemical library, the Medicines for Malaria Venture (MMV) Pathogen Box. This library was supplied as 1.0 x 10^−2^ M solutions in DMSO and was further diluted in DMSO to generate a 1.0 x 10^−3^ M stock of each compound. The library was then screened against An. gambiae using the INVAPP / Paragon system; all chemicals from the collection were initially tested at 1.0 x 10^−5^ M, n = 3, 1% v/v final DMSO. The actions of each chemical on the movement index was recorded and ranked by their median (see supplementary data Table S1), and the 5 compounds that most reduced the motility index (MMV007803, MMV637229, MMV687775, MMV687776, MMV688934) were used in further analysis. All 5 were re-sourced in powder form as a kind gift of MMV, prepared as a fresh DMSO stock, and tested for concentration-dependent actions over the range 3×10^−7^ – 1×10^−4^ M for both the G3 and Tiassale 13. Of these compounds, MMV688934 (tolfenpyrad) showed a marked concentration-dependent immobilization of larvae (Fig. 2). Tolfenpyrad is an inhibitor of complex I of the respiratory electron transport chain in the mitochondria. It has been successfully trialled in toxic sugar baits to kill mosquitoes [20]. A weaker action was observed for MMV637229 (Clemastine, a first-generation antihistamine for the treatment of hayfever) which has also shown to be effective in killing *Trypanosoma cruzi* and *Plasmodium falciparum* [21,22].

**Figure 2.**
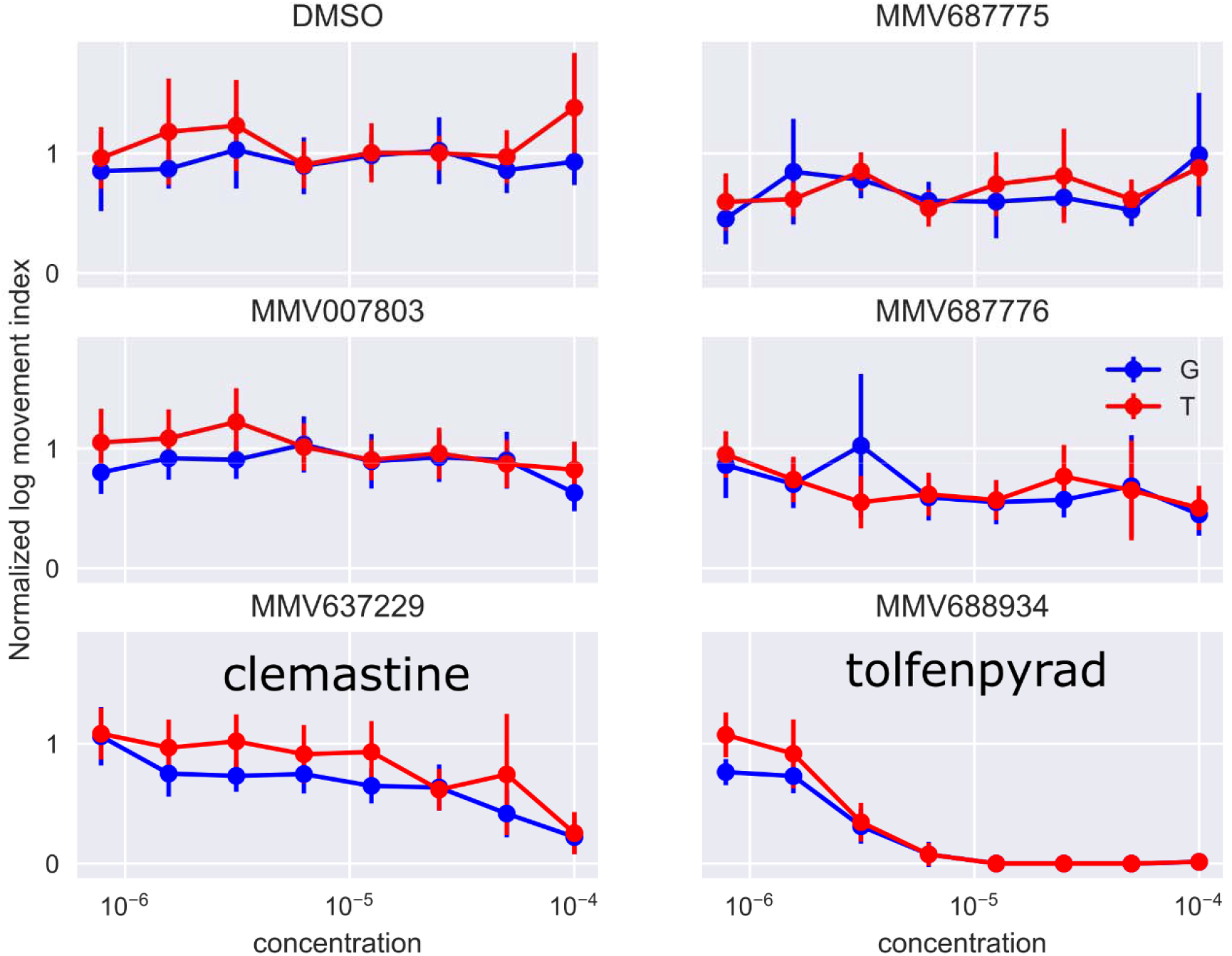
Screening a chemical library on *An. gambiae* larvae using the INVAPP / Paragon system. Concentration response curves for the five compounds in the Medicines for Malaria Venture (MMV) Pathogen Box library identified as active in the primary screen. The movement index for each well was measured before adding the compounds, and then again 240 min later. The second reading is divided by the first to normalize for background variations between the wells caused, for example, by differences in the number of larvae dispensed in each well. The compounds were applied over the range 3.0 × 10^−9^ M - 1.0 × 10^−4^ M, although only concentrations above 7.8 × 10^−7^ M are shown for clarity. Larval progeny from both resistant *An. coluzzii* (Tiassale) and sensitive *An. gambiae* (G3) strains were studied and labelled “G” and “T” in the figure respectively. Of the compounds tested, only MMV688934 (tolfenpyrad) and MMV637229 (Clemastine) have chemical names.

### Detecting motility differences in larval progeny from resistant and sensitive An. coluzzii and Ae. aegypti

We tested whether our INVAPP / Paragon system was able to detect behavioural differences in larvae derived from resistant compared to susceptible strains. Larvae from deltamethrin susceptible and resistance strains were examined for each species (G3 [susceptible] and Tiassale 13 [resistant] for Anopheles, and New Orleans [susceptible] and Cayman [resistant] for *Ae. aegypti*). Image sequences were filmed immediately before addition of Deltamethrin and at intervals (2, 5, 10, 20, 45 and 60 min) afterwards.

There was a concentration-dependent action of deltamethrin on motility for the 4 different strains (Fig. 3). The concentration dependence was detected after 5 min of exposure. At briefer exposures larvae were not completely immobilized, whereas at exposures longer than 60 min even the lowest concentrations tested had immobilized all larvae. The curves for the 4 strains after 60 min exposure to deltamethrin were fitted to a sigmoid function (see Methods), from which PIC_50_s were estimated.

**Figure 3.**
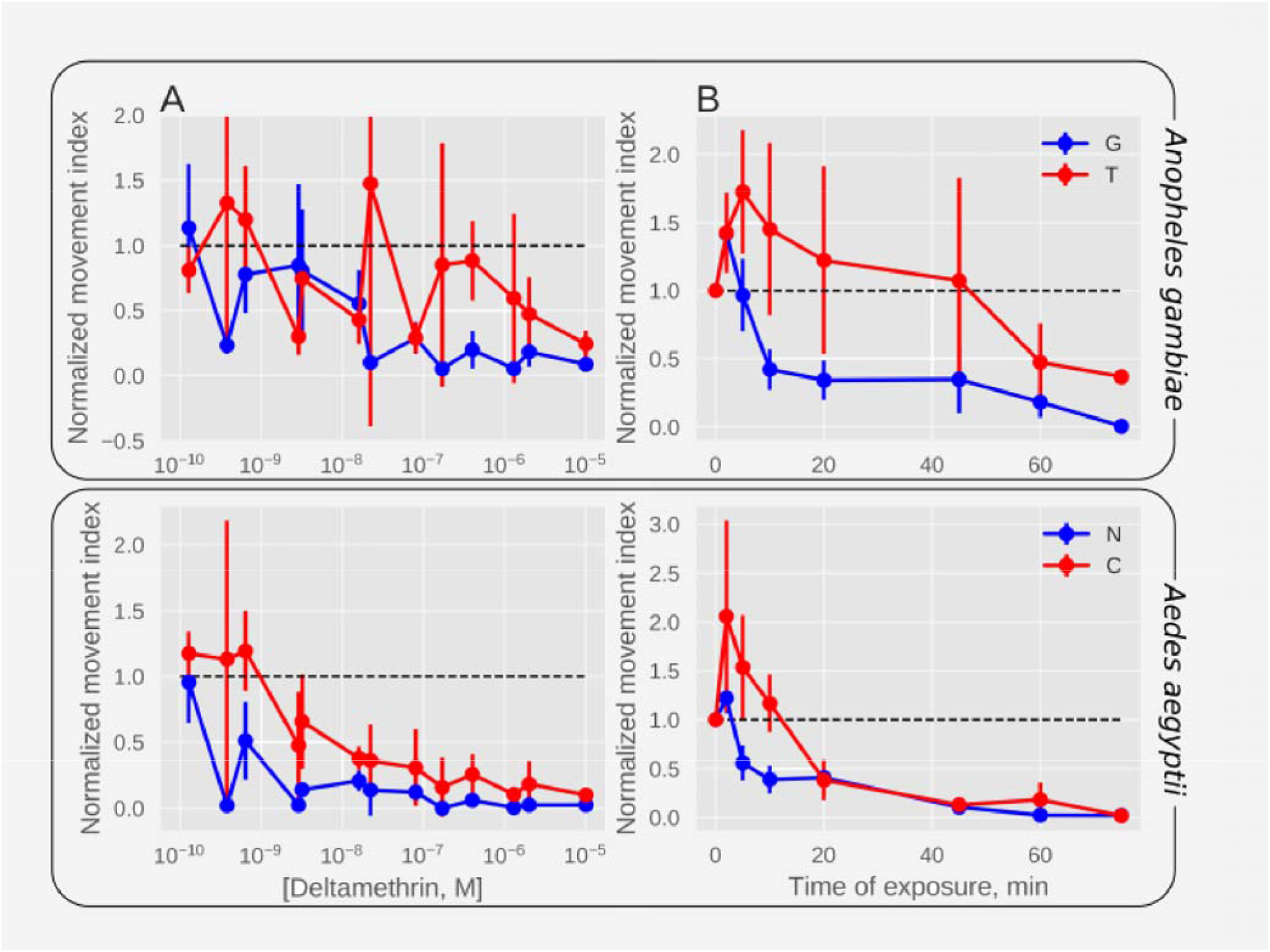
Motility measurement following deltamethrin exposure in concentration- and time-dependent manner for *An. Gambiae* susceptible (G3) and resistant (Tiassale) strains as well as for *Ae. Aegypti* susceptible (New Orleans) and resistant (Cayman) strains. The normalized movement index, which is the movement index (see Methods) divided by the index recorded before the application of insecticide, is plotted against the concentration after 60 min applications (A) and the time following exposure (B) to 10^−6^ M deltamethrin. Increased rate of movement inhibition is apparent in the larvae from adult susceptible strains (blue: G3 (G), New Orleans (N)) as compared to the adult resistant strains (red: Tiassale (T), Cayman (C)). The dotted line indicates the value before insecticide application. Error bars indicate ± **1** s.e.m.

The PIC_50_ for Tiassale was estimated to be 7.0 ± 1.0 (100 nM), while that for G3 was estimated to be 8.0 ± 1.0 (10 nM). Similarly, the PIC_50_ for Cayman was estimated to be 8.0 ± 1.0 (10 nM), while that for New Orleans was estimated to be 9.0 ± 0.0 (1 nM). The differences between susceptible and resistant strains were statistically significant (2-way ANOVA, F(1.0,24.0) = 7.0, P = 0.016) but no significant difference was detected between genus (2-way ANOVA, F(1.0,24.0) = 3.0, P = 0.091) and there was no statistically significant interaction between genus and susceptibility (2-way ANOVA, F(1.0,24.0) = 0.0, P = 0.906). Hence, the algorithm detected a more rapid decline in movement in larvae derived from resistant adults after exposure to deltamethrin in both Anopheles and Aedes species.

### Comparison of INVAPP / Paragon data with the standard WHO assay

To determine how the Paragon algorithm compared to the WHO larval assay, we tested the *Ae. aegypti* New Orleans and Cayman strains for deltamethrin resistance with traditional larval cup bioassays (Fig. 4). The WHO assay yielded logLC_50_s of 8.60 ± 0.04 (LC50 = 2.5 nM) and 7.29 ± 0.14 (LC50 = 51.3 nM) for New Orleans and Cayman respectively (2-tailed, independent t-test, T(4)=−3.9, P=0.0). The inhibition and resistance ratios estimated from the two methods by dividing the respective LC50s. Hence, the resistance ratio estimated from WHO assays is similar to the inhibition ratio obtained from Paragon. This suggests that the inhibition of movement at the 4hr time point in the Paragon assay correlates well with the 24hr killing observed in WHO assays.

**Figure 4.**
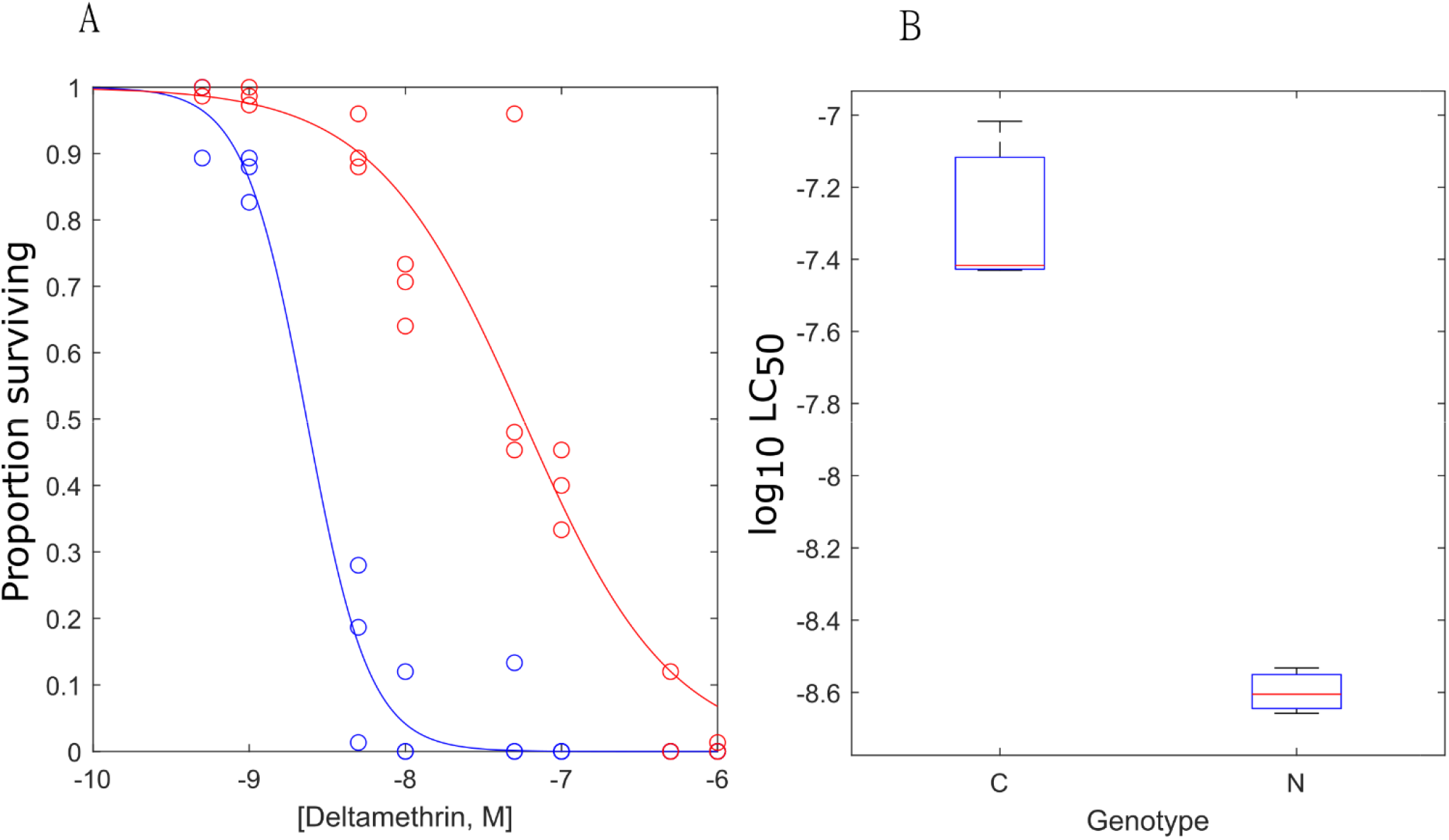
The application of the standard WHO larval assay to detect deltamethrin resistance in *Ae. aegypti*. Deltamethrin was tested over a range between 5.0 x 10^−10^ M to 1.0 x 10^−6^ M. (A) The concentration-response curves are given for the wild-type New Orleans strain (blue) and the Deltamethrin-resistant Cayman strain (red). (B) The fitted PIC_50_s estimated from fitting the curves in A to a Sigmoid curve. The WHO assay yielded LC_50_s of 8.60 ± 0.04 (2.5 nM) and 7.29 ± 0.14 (51.3 nM) for New Orleans and Cayman respectively (2-tailed, independent t-test, T(4)=−3.9, P=0.0). Each point in (B) represents the mean of 3 (WHO) separate experiments and error bars indicate the standard error of the mean.

### Smartphone detection of motility differences between pyrethroid susceptible and resistant strains

Larvae were dispensed into a petri dish or 96 well plate and allowed to settle. Films of 2000-3000 s duration were made at the smartphone camera default settings (LG G4: 16 MP, f/1.8, 28 mm). The INVAPP/Paragon algorithm was then applied with no changes other than the threshold for allocating a pixel as “having movement” or “not having movement”. In the absence of deltamethrin both larval strains yielded a roughly constant movement index over the duration of the recording (Fig. 5A, B, dotted lines). In the presence of 1 × 10^−6^ M deltamethrin, the movement index began to fall after about 500-1000 s to a steady-state, low level. The rate of this decline was slower for larvae from resistant strains (τ = 0.18 ± 0.08 x 10^3^ s^−1^ for Cayman and 3.5 ± 0.16 x 10^3^ s^−1^ for Tiassale) than it was for larvae from susceptible strains (τ = 2.7 ± 0.2 x 10^3^ s^−1^ for New Orleans and 12 ± 2.3 x 10^3^ s^−1^ for G3 – *Anopheles*: one-tailed unpaired t-test T(4) = 7.0, P = 0.003; *Ae. aegypti*: one-tailed unpaired t-test T(4) = 20.0, P < 0.001). Thus, INVAPP/Paragon detected the differential response of the larval progeny from resistant strains for two significant human disease vectors following short term exposure to 1 μM deltamethrin in real-time on a readily available smartphone.

**Figure 5.**
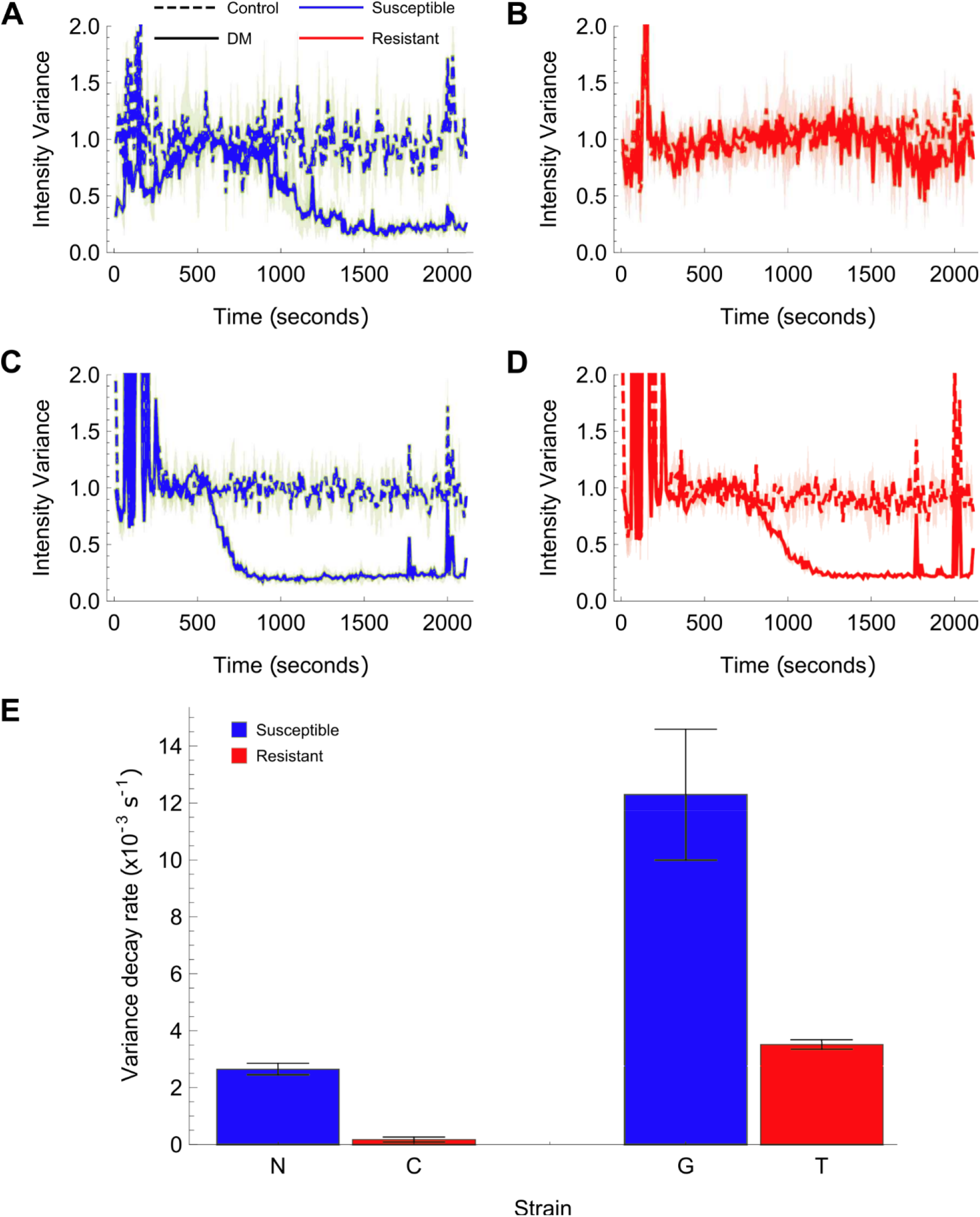
The output of a smartphone application that measures mosquito larvae motility following deltamethrin treatment, quantified by the average pixel variance over 2 s. (A) and (B) show the pixel variance over time (motility) of *Aedes aegypti* New Orleans and Cayman larvae with 10^−5^ M deltamethrin in water and DMSO/water only controls. The spike at t 200s is the addition of the deltamethrin or control (water). The solid lines show the mean and the shaded area, the standard deviation (n = 3). The susceptible strain (A) motility decreases over time with DM added, whilst the resistant strain (B) does not. (C) and (D) show the pixel variance over time of *Anopheles gambiae* G3 and *An. coluzzii* Tiassale larvae with 10^−5^ M deltamethrin in water and DMSO/water controls. In this case, the resistant strain variance also decreases, but at a slower rate. (E) Fitting the variance over time of all the DM samples to exponential decay functions allows the extraction of the decay rate in each case (see methods). These are plotted on the bar chart with error bars showing the standard deviation of the fitted decay rate (n = 3). A higher decay rate is observed for the larvae from susceptible strains compared to their respective resistant strains. *Aedes aegypti* p = 1.9 × 10^−5^ (one-tailed t-test, T(2)=19.8); *Anopheles* p = 1.1 × 10^−2^ (one-tailed t-test, T(2)=6.6).

## Discussion

Here we evaluate the utility of an automated screening platform / algorithm combination in quantifying the actions of chemicals tested against mosquito larvae for their capacity to impair larval motility. Previously, the “<u>I</u>N<u>V</u>ertebrate <u>A</u>utomated <u>P</u>henotyping platform (INVAPP), with the Paragon algorithm [13] had been shown to enable high-throughput chemical screening on the nematode genetic model organism C. elegans and parasitic nematodes such as *Haemonchus contortus, Teladorsagia circumcincta*, and *Trichuris muris* worms in the search for novel anthelmintic candidates [13]. The system allows 96 well plate-based chemical screening for anthelmintic activity. By this means we detected compounds affecting the motility and development of worms. We deployed known anthelmintics to validate the utility of the INVAPP/Paragon system and screened the MMV Pathogen Box chemical library identifying compounds already known to have anthelmintic or anti-parasitic activity and other compounds previously not known to have anthelmintic activity [13,23]. In separate studies we showed how this approach could help identify novel classes of chemistry with anthelmintic activity, the dihydrobenz[e] [1,4] oxazepin-2 *(3H)*-ones [24,25] and the 2,4-diaminothienol [3,2-*d*] pyrimidines [26].

Here we demonstrate the use of INVAPP paired with the analysis algorithm Paragon as a rapid, convenient and robust method for estimating a biologically-relevant parameter (motility) of the behaviour of mosquito larvae from two key human disease vectors. It was first used to detect the actions on motility of *Ae. aegypti* larvae of a known larvicide, temephos. Following this validation of the ability of INVAPP / Paragon to detect larval motility impairment by an established larvicide, we explored its potential use in library-scale chemical screening. The discovery of new and re-profiled insecticidally-active chemicals remains important. As a proof-of-principle exercise, we screened (blind) a 400-compound library – The Medicines for Malaria Venture (MMV) Pathogen Box. We identified tolfenpyrad, an established insecticide, as a toxic agent for mosquito larvae. Tolfenpyrad is a pyrazole insecticide developed by the Mitsubishi Chemical Corporation in 1991. It is active against adults of the Anopheles strains used in this study (Lees et al 2020), whiteflies [27] and aphids [28] as well as *H. contortus* [29]. Since tolfenpyrad is the only known insecticide in the MMV Pathogen Box, this is indicative of the specificity of the screen.

As well as readily identifying the potent insecticide, tolfenpyrad, our screen of the MMV Pathogen Box also showed that clemastine has a paralysing action on mosquito larvae, suggesting that the algorithm can be useful in high throughput chemical re-profiling screens. Clemastine is a sedating anti-histaminergic drug used in the treatment of allergic reactions, including seasonal allergic rhinitis (“hay-fever”) [30]. In vertebrates it blocks primarily the H1-subytpe of histamine receptor, which is a member of the 7-transmembrane, G-protein-coupled receptor family [31]. Insects are known to use histamine as a neurotransmitter and to possess a histamine-gated chloride channel with some pharmacological similarities to the vertebrate H1 class [31,32]. Perhaps of equal interest is the fact that histamine-gated chloride channels are not present in vertebrates [33] and insects do not possess a metabotropic histamine receptor [34]. Cys-loop ligand-gated anion channels are known targets of insecticides and anthelmintics [35,36]. It has also been reported that mutations in the two Drosophila histamine-gated chloride channels (HClA and HClB) have opposite effects on ivermectin sensitivity: mutation of HClA confers enhanced sensitivity whereas knockout of HClB diminished sensitivity to avermectin neurotoxins [37,38], suggesting that these channels as well as GluCls may play a role in the action of macrocylic lactone derivatives. Furthermore, the finding that two histamine-gated chloride channels cloned from the housefly, *Musca domestica*, are insensitive to other insecticides known to target GABA and glutamate-gated ion channels [39] suggests that histamine-gated chloride channels may have a very distinct pharmacological profile offering a quite new candidate insecticidal target. However, it should be noted that neither HClA [40] nor HClB [41] null mutants are lethal, suggesting a possible limitation on the usefulness of these receptors as targets of insecticides.

Resistance to drugs and chemicals used to combat malaria seriously threatens the success of control measures and the investment therein over the past 15 years. The widespread emergence of pyrethroid resistance threatens the hitherto successful deployment of ITNs [3,41]. Resistance in the adult mosquito vectors is established but much less is known concerning resistance in mosquito larvae for which targeting by larvicides is a valid control approach especially for *Ae. aegypti*. Here using behavioural phenotyping, we have shown that comparison of susceptible and resistant *Ae. aegypti* larvae, produced a ratio of reduction in movement index that was similar to the resistance ratio determined through WHO assays. A comparative reduction in motility was also observed following exposure of Anopheles larvae from strains where the adults show WHO resistant phenotypes. These findings are robust in our laboratory studies, but will require the analysis of more resistant strains to determine the precise predictive nature of INVAPP / Paragon measurements compared to WHO classified resistance ratio. In future, it will also be of interest to explore whether dual larvicidal treatments can help combat resistance and whether the INVAPP / Paragon system can be used to assist in identifying new areas of chemistry and / or new targets not already compromised by known resistance mechanisms that will be needed for such approaches.

INVAPP combined with the Paragon algorithm would allow for faster measurement than can be accomplished using the currently deployed manual WHO assay for larvicidal activity. It may not be as sensitive as the manual assay, but it allows the effectiveness of compounds to be measured more rapidly, and provides more detail in the time-dependence of the actions of compounds to be identified. It also requires a smaller bench footprint and is less labour intensive. Also, manual assays such as the current WHO assay are subject to investigator fatigue and the end-point assessment is not always straightforward. INVAPP / Paragon, which is automated, reduces errors due to these factors.

Increasingly, sensitive smartphone applications and cameras are being used for the diagnosis and study of infectious diseases including malaria. For example, a smartphone polarised microscope has been used for malaria diagnosis [42]. Automated detection of malaria parasites in blood smears has been achieved [43,44]. A rapid and robust field detection of resistance that utilises a simple smartphone assay could also yield valuable information. Geo-located data on larval resistance could be readily transmitted to central databases that provide regional information of the temporal and spatial development of insecticide resistance which can be mapped alongside the emergence of adult resistance and malaria prevalence data [45,46]. We have shown potential to detect differences in behavioural response to a pyrethroid insecticide using a smartphone camera. This paves the way for the development of a smartphone application perhaps to be used in conjunction with a simple, slide-on test kit that will facilitate such measurements. These assays are a promising first step in developing such a simple-to-operate field assay.

## Acknowledgements

We thank Medicines for Malaria Venture for the Pathogen Box library and resupply of hit compounds.

---

**Supplementary Table S1.** The top 20 hits identified in a screen of the Malaria for Medicines Venture box. The rows are sorted by the values of the movement index. The phenotype and location in the box are shown. In the “Phenotype” column, G stands for G3 (susceptible) and T for Tiassale (resistant).

**Supplementary Table S2.** The raw data from the Malaria for Medicines Venture box are shown, with details provided with the MMV box. In the “Phenotype” column, G stands for G3 (susceptible) and T for Tiassale (resistant).

