## Supplemental Table S1 for "Automated phenotyping of mosquito larvae enables high-throughput screening for novel larvicides and offers potential for smartphone-based detection of larval insecticide resistance"

| phenotype | box | row | column | Movement index |
| --- | --- | --- | --- | --- |
| G | B | 4 | 4 | 32.46667 |
| G | B | 1 | 4 | 41.26667 |
| G | B | 5 | 4 | 45 |
| G | D | 6 | 1 | 55.73333 |
| G | A | 1 | 5 | 56.26667 |
| T | D | 3 | 10 | 57.6 |
| G | E | 0 | 1 | 59.1 |
| T | D | 7 | 3 | 59.7 |
| T | D | 0 | 1 | 60.1 |
| T | C | 7 | 11 | 60.5 |
| G | B | 1 | 2 | 63.8 |
| T | D | 0 | 7 | 66.7 |
| G | C | 7 | 9 | 68.8 |
| T | D | 7 | 2 | 69.2 |
| G | D | 7 | 9 | 72.33333 |
| G | B | 7 | 4 | 74 |
| G | C | 7 | 4 | 74.06667 |
| T | D | 0 | 4 | 74.9 |
| G | B | 2 | 1 | 76.8 |
| T | D | 6 | 3 | 76.9 |
